# SETDB1 Triple Tudor Domain Ligand, (*R,R*)-59, Promotes Methylation of Akt1 in Cells

**DOI:** 10.1101/2023.05.10.539986

**Authors:** Mélanie Uguen, Yu Deng, Fengling Li, Devan J. Shell, Jacqueline L. Norris-Drouin, Michael A. Stashko, Suzanne Ackloo, Cheryl H. Arrowsmith, Lindsey I. James, Pengda Liu, Kenneth H. Pearce, Stephen V. Frye

## Abstract

Increased expression and hyperactivation of the methyltransferase SETDB1 are commonly observed in cancer and central nervous system disorders. However, there are currently no reported SETDB1-specific methyltransferase inhibitors in the literature, suggesting this is a challenging target. Here, we disclose that the previously reported small-molecule ligand for SETDB1’s Triple Tudor Domain, (*R,R*)-59, is unexpectedly able to increase SETDB1 methyltransferase activity both *in vitro* and in cells. Specifically, (*R,R*)-59 promotes *in vitro* SETDB1-mediated methylation of lysine 64 of the protein kinase Akt1. Treatment with (*R,R*)-59 also increased Akt1 threonine 308 phosphorylation and activation, a known consequence of Akt1 methylation, resulting in stimulated cell proliferation in a dose-dependent manner. (*R,R*)-59 is the first SETDB1 small-molecule positive activator for the methyltransferase activity of this protein. Mechanism of action studies show that full-length SETDB1 is required for significant *in vitro* methylation of an Akt1-K64 peptide, and that this activity is stimulated by (*R,R*)-59 primarily through an increase in catalytic activity rather than a change in SAM binding.

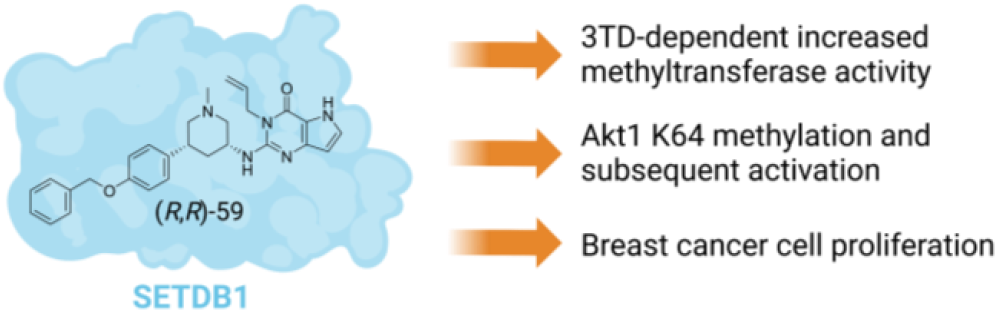

## INTRODUCTION

The lysine methyltransferase protein class is mostly known for its ability to methylate lysine residues of histone proteins and modulate chromatin accessibility and transcription. This protein class contains two subsets including 55 SET domain-containing methyltransferases and 131 7βS domain-containing methyltransferases.^1^ In addition to histone methylation, many of these methyltransferases are able to install methyl groups on lysines from non-histone proteins which have been linked to activity regulation.^2–4^

Among these methyltransferases, SET domain bifurcated 1 (SETDB1) has been shown to methylate non-histone proteins, such as the protein kinase, Akt1, at residues K64, K140 and K142, and p53 at residue K370. SETDB1-mediated methylation of both Akt1 and p53 are associated with tumorigenesis.^5–7^ However, SETDB1 is mostly known for its histone methylation properties as it is the only lysine methyltransferase known to di- and trimethylate lysine 9 of histone 3 (H3K9) in euchromatic regions resulting in gene repression.^8^ Methylation is mediated by the catalytic SET domain of SETDB1 through transfer of a methyl group from the S-adenosyl methionine (SAM) cofactor to the substrate.^9^ Additionally, SETDB1 contains other structured domains including a methyl-CpG DNA binding domain and a Triple Tudor Domain (3TD) (Figure 1).^8^ The 3TD is a chromatin reader domain which binds the histone 3 tail when methylated at lysine 9 and acetylated at lysine 14, with highest affinity for K9 dimethylation.^10^

**Figure 1.**
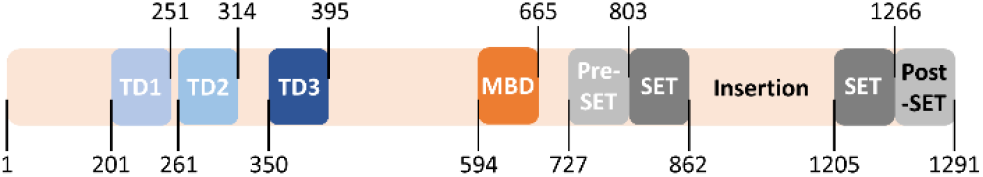
SETDB1 contains a Triple Tudor Domain, which includes Tudor domain 1 (TD1), Tudor domain 2 (TD2) and Tudor domain 3 (TD3), a Methyl-CpG binding domain (MBD) and the catalytic domain made of a Pre-SET, two SET and a Post-SET domain.

Generally, SETDB1 overexpression and aberrant hyperactivation are widely observed in many cancer types such as breast cancer, prostate cancer, non-small cell lung cancer, colorectal cancer, glioma, melanoma, and others, and are associated with poor prognsis.^11^ These findings demonstrate the potential of SETDB1 inhibition as a promising anti-cancer treatment. Aside from cancer, SETDB1 overexpression has also been linked with central nervous system disorders such as Huntington’s disease, schizophrenia, and autism, widening the potential applications of SETDB1-targeting therapeutics.^12^ However, there are currently no available inhibitor that are able to selectively modulate the methyltransferase activity of SETDB1.

During our investigation of small molecule SETDB1 ligands, we discovered the unexpected ability of a recently reported TD2 ligand, (*R,R*)-59, to increase the methyltransferase activity of the SET domain both *in vitro* and in cells. *In vitro*, (*R,R*)-59 increased the amount of methylated Akt1-derived peptide in a radioactivity-based methyltransferase assay. This activation was also observed in cells where an increase in Akt1 trimethylation and resulting phosphorylation, which is known to be a SETDB1-mediated process, were observed. As described below, we show that (*R,R*)-59 is the first SETDB1 ligand for positive modulation of its methyltransferase activity.

## RESULTS AND DISCUSSION

### SETDB1 3TD is Required for the Methylation of Akt1 K64

Akt1 is believed to be uniquely methylated by SETDB1 in contrast to histone H3, which is a substrate for other methyltransferases, such as Suv39H1, Suv39H2, G9A and GLP.^1^ Therefore, we focused on Akt1 as a substrate and started by examining the ability of SETDB1 to methylate Akt1 *in vitro* using a tritiated SAM-based methyltransferase activity assay. Briefly, active recombinant SETDB1 protein was incubated with a biotinylated Akt1 peptide and ^3^H-SAM. A biotin binding membrane allowed for isolation of the peptide, and the levels of ^3^H were quantified and proportional to the activity of the protein. Two Akt1-derived peptides were tested as methyltransferase substrates: Akt1-K64 (aa 59-80), containing the previously reported K64 trimethylation site,^6^ and Akt1-K140 (aa 131-152), containing two other reported Akt1 trimethylation sites, K140 and K142.^7^ Results showed that Akt1-K64 is a substrate for full-length SETDB1 (SETDB1-FL) with an observed increase in activity with increasing protein concentrations (Figure 2). However, significantly less methyltransferase activity was observed with the Akt1-K140 peptide compared to Akt1-K64 (Figure 2). These results are in agreement with the findings from Wang *et al*. who detected K64 as the major SETDB1-mediated Akt1 methylation site by mass spectrometry analysis.^6^ Although Guo *et al*. were able to observe *in vitro* Akt1 methylation of K140/142 by SETDB1, it is possible that the levels were too low to be detected by our assay.^7^

**Figure 2.**
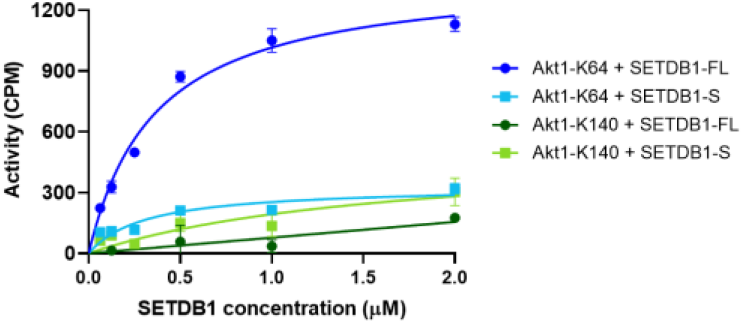
Evaluation of Akt1-derived peptides as substrates for the SETDB1 methyltransferase activity assay. We attempted to methylate the full-length Akt1 protein *in vitro* but no significant methylation was observed in presence of 1 μM of Akt1. It is possible that the methylation of Akt1-K64 requires attachment of Akt1 PH domain to PIP3 to expose K64 residue for SETDB1 recognition^13^, or the Akt1 concentration was not high enough to observe significant activity in this assay.

Finally, to evaluate the influence of the 3TD on the methyltransferase activity of SETDB1, we examined the ability of a shorter SETDB1 construct (aa 590-1291), SETDB1-S, to induce Akt1-K64 methylation *in vitro*. This shorter construct contains all ordered domains except the 3TD and is still catalytically active towards H3. Results showed no significant methylation of Akt1-K64 by SETDB1-S as there was again only limited increase in ^3^H levels compared to SETDB1-FL (Figure 2). This suggests that the 3TD of SETDB1 is required for the efficient methylation of K64 by Akt1. We demonstrated that full-length SETDB1 is able to methylate Akt1 K64 but not K140 nor K142 and K64 methylation likely requires the presence of the 3TD as the SETDB1-S construct was not able to methylate Akt1 K64 *in vitro*.

### SETDB1-mediated Methylation is Promoted by (*R,R*)-59 Binding

To study the relationship between the SETDB1 3TD reader domain and the catalytic SET domain, we examined the influence of (*R,R*)-59, a potent small-molecule that binds the aromatic cage of the TD2, on SETDB1 catalytic activity (Figure 3). We first characterized the binding affinity of the compound to the isolated 3TD. (*R,R*)-59 was synthesized following the procedure described by Guo *et al*.^14^ (*R,R*)-59 was tested in a competition time-resolved fluorescence energy transfer (TR-FRET) assay providing an IC50 of 1.2 ± 0.14 μM for the displacement of the H3K9Me2K14Ac (1-20) peptide (Figure S1). In a differential scanning fluorimetry (DSF) assay, the ΔTm was 2.9 ± 0.20 °C suggesting that (*R,R*)-59 is able to stabilize the 3TD of SETDB1. We tested the compound in a direct binding assay using surface plasmon resonance (SPR), which yielded a KD of 4.8 ± 1.6 μM (Figure S2). Using TR-FRET, we also confirmed that (S,S)-59 (Figure 3), the enantiomer of (R,R)-59, was not able to displace the H3K9Me2K14Ac (aa 1-20) peptide.

**Figure 3.**
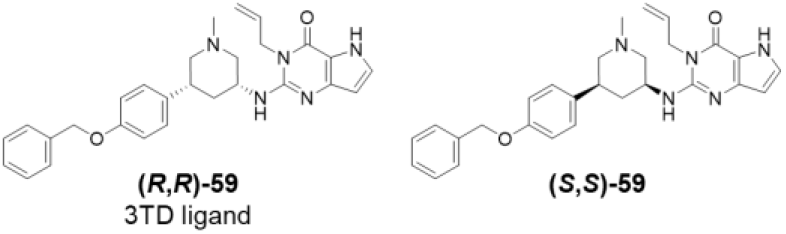
Structure of SETDB1 3TD ligand (*R,R*)-59 and enantiomer (*S,S*)-59.

Although our protein constructs and binding assays showed diminished affinity relative to prior reports,^14^ we confirmed that (*R,R*)-59 is a ligand for the 3TD of SETDB1. We then examined its ability to modulate the methyltransferase activity of SETDB1. Surprisingly, in the SETDB1-FL *in vitro* methyltransferase assay, (*R,R*)-59 was able to promote methylation of the Akt1-K64 (aa 59-80) peptide in a dose-dependent manner. We observed an increase in the activity of SETDB1-FL by up to 50% in the presence of 100 μM of (*R,R*)-59 (Figure 4A). Moreover, increasing concentrations of (*R,R*)-59 resulted in a dose-dependent increase in SETDB1 activity in a SAM-dependent manner (Figure 4B). Overall, this experiment shows that (*R,R*)-59 can promote the methyltransferase activity of SETDB1 at high concentrations *in vitro*.

**Figure 4.**
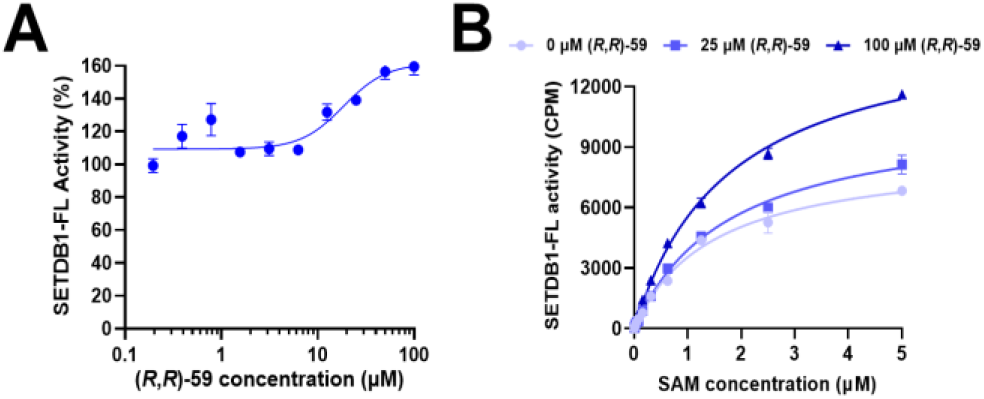
(A) Modulation of the methylation of an unmodified Akt1-K64 peptide as a function of the concentration of (*R,R*)-59. (B) Kinetic evaluation of the influence of SAM on the enzymatic activity of SETDB1 with and without (*R,R*)-59.

We then examined the influence of the small molecule on the kinetic parameters of the cofactor SAM in the presence of various concentrations of (*R,R*)-59 (Figure 4B). An increase in kcat was observed in the presence of (*R,R*)-59, but the KM for SAM remained constant (Table 1). The increase in the catalytic efficiency (kcat/KM) from 0.32 to 0.49 μM^-1^h^-1^ suggests an increased efficiency of the methyltransferase properties of SETDB1 in the presence of (*R,R*)-59, explaining the observed increase in the amount of methylated Akt1-K64. We attempted to evaluate the kinetic parameters KM and kcat for Akt1-K64 but we did not reach steady state when using concentrations up to 100 μM of the peptide substrate (Figure S3).

**Table 1.** Kinetic parameters for SAM in the presence of different concentrations of (*R,R*)-59.

|  | (R,R)-59 concentration |  |  |
| --- | --- | --- | --- |
| | 0 μM | 25 μM | 100 μM |
| SAM $K_M$ (μM) | $0.90 \pm 0.08$ | $0.92 \pm 0.03$ | $1.02 \pm 0.02$ |
| SAM $k_{cat}$ ($h^{-1}$) | $0.29 \pm 0.01$ | $0.34 \pm 0.02$ | $0.50 \pm 0.01$ |
| SAM $k_{cat}/K_M$ ($\mu M^{-1}h^{-1}$) | 0.32 | 0.37 | 0.49 |

Overall, we showed that (*R,R*)-59 is able to increase the SETDB1 methyltransferase activity *in vitro* by enhancing the efficiency of the methyl transfer from SAM to the Akt1-K64 peptide. This may be due to an allosteric interaction where (*R,R*)-59 is binding the TD2 domain to result in an enhanced methyltransferase activity of SETDB1 thanks to a more efficient SAM-mediated methyl transfer mechanism.

### (*R,R*)-59 Promotes Akt1 Trimethylation and Activation in Cells

Next, we proceeded to test the effect of (*R,R*)-59 in modulating methylation in cells. It was recently reported that SETDB1-mediated Akt1 methylation at K64, K140 and K142 residues promotes Akt1 phosphorylation at T308 and subsequent activation, contributing to oncogenicity and facilitating tumorigenesis.^6,7^ 24-hour treatment of HEK293T cells transfected with HA-tagged Akt1 treated with increasing concentrations of (*R,R*)-59 triggered Akt1 trimethylation and T308 phosphorylation in a dose-(Figure 5A) and time-dependent (Figure 5B) manner. Importantly, this effect was specific to (*R,R*)-59 as no increases of Akt1 trimethylation nor phosphorylation were observed after treatment with the non-binding enantiomer (*S,S*)-59 (Figure 5C). (*R,R*)-59 induced Akt1 methylation and phosphorylation were also observed in DLD1 cells stably expressing a lenti-viral HA-tagged Akt1 (Figure 5D).

**Figure 5.**
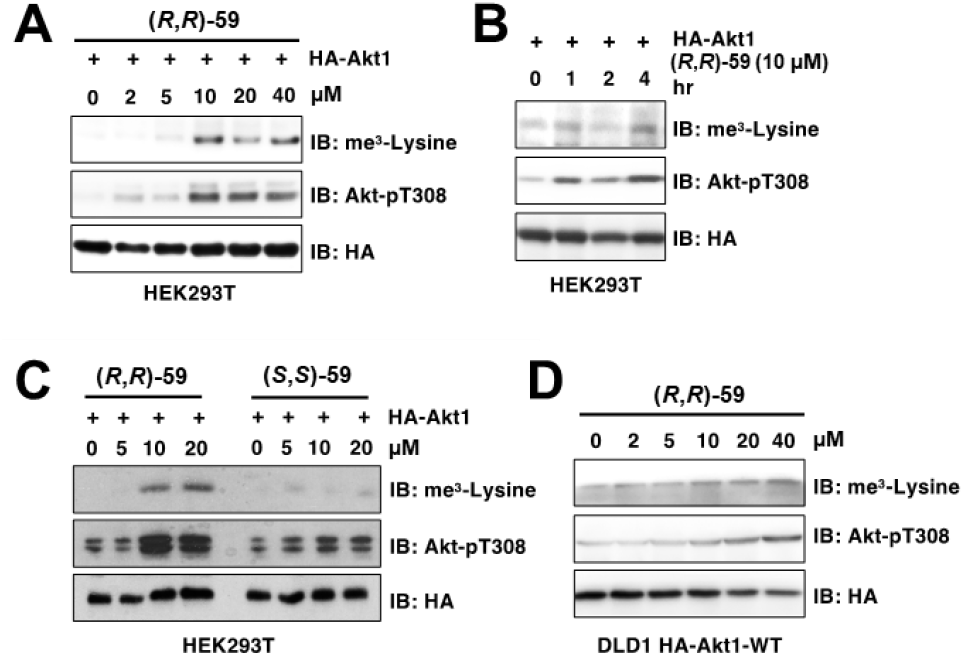
(*R,R*)-59 increases Akt1 methylation and subsequent phosphorylation after cell treatment. HEK293T cells transfected with HA-Akt1 showed increased trimethylation and T308 phosphorylation after 24-hour treatment with (*R,R*)-59 in a dose-(A) and time-dependent (B) manner. (C) (*S,S*)-59 showed no modulation of SETDB1 activity in cells. (D) A dose-dependent increase in methylated and phosphorylated Akt1 was also observed in DLD1 cells expressing a lenti-viral Akt1.

Querying TCGA datasets showed that the *SETDB1* gene is amplified in breast cancer (Figure 6A). Thus, we further examined the effects of (*R,R*)-59 on breast cancer proliferation, given that Akt1 hyperactivation has been observed in breast cancer and is thought to be critical to growth and drug resistance.^15^ We found that treatment of MDA-MB-231 cells with lower doses (2.5 μM and 5 μM) of (*R,R*)-59 but not its negative control compound promoted cell proliferation (Figure 6B); however, at higher dosed (> 5 μM) this effect is no longer observed. Together, these data suggest that (*R,R*)-59 exerts potent in-cell activities in modulating cell proliferation through promoting Akt1 methylation and activation.

**Figure 6.**
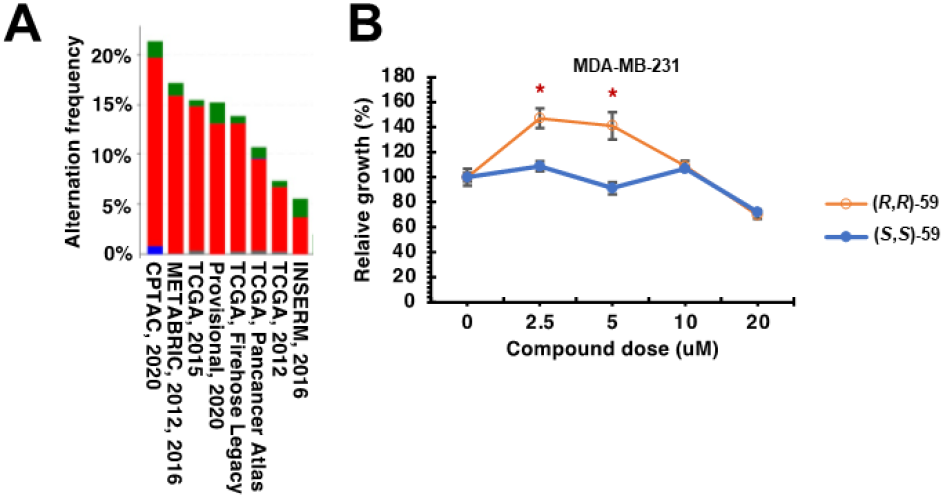
SETDB1 hyperactivation promotes triple negative breast cancer cell proliferation. (A) TGCA query for SETDB1 gene amplification. (B) (*R,R*)-59 promotes cell proliferation in triple negative breast cancer cells (MDA-MB-231) at specific concentrations while the negative control (*S,S*)-59 has no effect.

## MATERIALS AND METHODS

### Protein and peptide production and purification

A pET28-mhl vector containing the coding region for the SETDB1 Triple Tudor domain (residues 196-403 of NP_036564) was transformed into Rosetta2 BL21(DE3)pLysS competent cells (Novagen, EMD Chemicals, San Diego, CA). A 6 L culture was grown to mid log phase at 37 °C at which time the temperature was lowered to 18 °C and protein expression was induced by addition of 0.5 mM IPTG. Expression was allowed to continue overnight. Cells were harvested by centrifugation and pellets were stored at -80°C.

SETDB1 Triple Tudor protein was purified by resuspending thawed cell pellets in 30 mL of lysis buffer (50 mM sodium phosphate pH 7.2, 50 mM NaCl, 30 mM imidazole, 1X EDTA free protease inhibitor cocktail (Roche Diagnostics, Indianapolis, IN)) per liter of culture. Cells were lysed on ice by sonication with a Branson Digital 450 Sonifier (Branson Ultrasonics, Danbury, CT) at 40% amplitude for 12 cycles with each cycle consisting of a 20 second pulse followed by a 40 second rest. The cell lysate was clarified by centrifugation and loaded onto a HisTrap FF column (Cytiva, Marlborough, MA) that had been preequilibrated with 10 column volumes of binding buffer (50 mM sodium phosphate pH 7.2, 500 mM NaCl, 30 mM imidazole) using an AKTA FPLC (Cytiva, Marlborough, MA). The column was washed with 15 column volumes of binding buffer and protein was eluted in a linear gradient to 100% elution buffer (50 mM sodium phosphate pH 7.2, 500 mM NaCl, 500 mM imidazole) over 20 column volumes. Peak fractions containing SETDB1 Triple Tudor protein were pooled and concentrated to 2 mL in Amicon Ultra-15 concentrators 10,000 molecular weight cut-off (Merck Millipore, Carrigtwohill Co., Cork, Ireland). Concentrated protein was loaded onto a HiLoad 26/60 Superdex 75 prep grade column (Cytiva, Marlborough, MA) that had been preequilibrated with 1.2 column volumes of sizing buffer (25 mM Tris pH 7.5, 250 mM NaCl, 5% glycerol, 2 mM DTT) using an ATKA Purifier (Cytiva, Marlborough, MA). Protein was eluted isocratically in sizing buffer over 1.3 column volumes at a flow rate of 2 mL/min collecting 3 mL fractions. Peak fractions were analyzed for purity by SDS-PAGE and those containing pure protein were pooled and concentrated using Amicon Ultra-15 concentrators 10,000 molecular weight cut-off (Merck Millipore, Carrigtwohill Co. Cork TRL). Concentrated proteins were aliquoted and stored at -80°C in sizing buffer.

The truncated SETDB1 (570-1291aa, SETDB1-S) cDNA was cloned into the vector pFBOH-SUMOstar-TEV, and the full-length SETDB1 (2-1291aa, SETDB1-FL) cDNA was cloned into the vector pFastBac HTA. Both were expressed in sf9 insect cells and purified sequentially using Ni-NTA affinity (for SETDB1-S) or anti-FLAG affinity (for SETDB1-FL), and size exclusion chromatography to apparent homogeneity. Purified proteins were stored at - 80°C in storage buffer (50 mM Tris 7.4, 150 mM NaCl, 5mM DTT,10% glycerol) until use.

Akt1-K64 and Akt1-K140 peptides were purchased from Genscript. The countersalts are hydrochloric salts. The sequences are as followed:

Akt1-K64: QCQLMKTERPRPNTFIIRCLQW{K(Biotin)}, disul-fide bridge between C2 and C19 of the peptide.

Akt1-K140: AEEMEVSLAKPKHRVTMNEFEY{K(Biotin)}

The recombinant Akt1 protein (cat. No 81154) was purchased from Active Motif (Carlsbad, CA).

### Radiometric Methyltransferase Assay

The methyltransferase activities of full length-SETDB1 (2-1291aa) and truncated SETDB1 (570-1292aa) were measured using a radiometric assay that detects transfer of a [3H] methyl from the radio-labeled cofactor S-[Methyl-3H]-adenosyl-L-methionine (SAM, PerkinElmer, cat. NET155001MC) to a biotinylated peptide Akt1-K64 (QCQLMKTERPRPNTFIIRCLQW) or Akt1-K140 (AEEMEVSLAKPKHRVTMNEFEY) substrate. The biotinylated peptides were synthesized by GenScript (Piscataway, NJ, USA).

10 μM of biotinylated Akt1-K64 or Akt1-K140 as substrates were evaluated with 5 μM of 3H-SAM and varying concentrations of SETDB1-FL or SETDB1-S (up to 2 μM).

The kinetics of SAM on SETDB1-FL activity was assessed in the presence or absence of (*R,R*)-59 (0 μM, 25 μM and 100 μM) at 0.5 μM of SETDB1-FL, 100 μM of biotinylated Akt1-K64 and varying concentrations of 3H-SAM (up to 5 μM).

The effect of small-molecule (*R,R*)-59 on SETDB1-FL activity was determined at 0.5 μM of SETDB1-FL, 10 μM of biotinylated Akt1-K64 and 5 μM of 3H-SAM.

All enzymatic reactions (in triplicate) were performed at 23 °C with 60 min incubation in buffer consisting of 20 mM Tris (pH 8.5), 5 mM DTT, 0.01% Triton X-100. To stop the reactions, 10 μL of 7.5 M guanidine hydrochloride was added to 10 μL of reaction mixture and incorporated radioactivity was captured on SAM2® Biotin Capture Membranes (Promega, Madison, WI) and quantified using liquid scintillation counting as described before.^16^

### Ligand preparation

#### General information

Chemicals were purchased from commercial suppliers and used without further purification. Thin layer chromatography (TLC) was performed on glass plates coated with 60 F254 silica. Flash chromatography was carried out using a Teledyne Isco Combiflash Rf200, Rf200i or NextGen 300+ automated flash system with RediSep Rf normal phase or C18 RediSep Rf Gold reverse phase silica gel pre-packed columns. Fractions were collected at 220 nm and/or 254 nm. Preparative HPLC was performed using an Agilent Prep 1200 series with the UV detector set to 220 nm and 254 nm. Samples were injected onto either a Phenomenex Luna 250 × 30 mm (5 μm) C18 column or a Phenomenex Luna 75 x 30 mm (5 μm) C18 column at room temperature.

#### Analytical equipment

^1^H NMR spectra were obtained using a Varian 400MR Inova spectrometer using a frequency of 400 MHz. ^13^C spectra were acquired using the Varian 400MR Inova spectrometer operating at a frequency of 101 MHz or a Bruker Avance III HD 700 MHz at a frequency of 176 MHz. The abbreviations for spin multiplicity are as follows: s = singlet; d = doublet; t = triplet; q = quartet, p = quintuplet, h = sextuplet and m = multiplet. Combinations of these abbreviations are employed to describe more complex splitting patterns (e.g. dd = doublet of doublets). Analytical LCMS data for all compounds was acquired using an Agilent 6110 Series system with the UV detector set to 254 nm. Samples were injected (<10 μL) onto an Agilent Eclipse Plus 4.6 × 50 mm, 1.8 μm, C18 column at room temperature. Mobile phases A (H2O + 0.1% acetic acid) and B (MeCN + 1% H2O + 0.1% acetic acid) were used with a linear gradient from 10% to 100% B in 5.0 min, followed by a flush at 100% B for another 2 minutes with a flow rate of 1.0 mL/min. Mass spectra (MS) data were acquired in positive ion mode using an Agilent 6110 single quadrupole mass spectrometer with an electrospray ionization (ESI) source. Analytical LCMS (at 254 nm) was used to establish the purity of targeted compounds. All compounds that were evaluated in biochemical and biophysical assays had >95% purity as determined by LCMS.

#### Compound data

##### (R,R)-59-3-allyl-2-{[(3R,5R)-5-(4-(benzyloxy)pheny)-1-methylpiperidin-3-yl]amino}-3,5-dihydro-4H-pyrrolo[3,2-d]pyrimidin-4-one

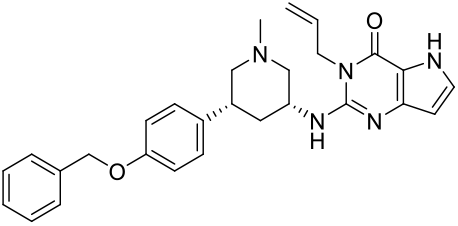

(*R,R*)-59 was prepared according to the procedure described in the literature.^14^ 6.7 mg of white solid were obtained. Rf (10% MeOH in DCM) = 0.71. ^1^H NMR (400 MHz, cd3od) δ 7.44 – 7.36 (m, 2H), 7.36 – 7.29 (m, 2H), 7.29 – 7.23 (m, 1H), 7.20 (d, J = 2.9 Hz, 1H), 7.18 – 7.09 (m, 2H), 6.95 – 6.86 (m, 2H), 6.12 (d, J = 2.8 Hz, 1H), 5.97 – 5.83 (m, 1H), 5.16 (dq, J = 10.6, 1.7 Hz, 1H), 5.07 – 4.97 (m, 3H), 4.78 – 4.70 (m, 2H), 4.38 – 4.25 (m, 1H), 3.23 (dd, J = 10.6, 4.0 Hz, 1H), 2.98 – 2.84 (m, 2H), 2.32 (s, 3H), 2.17 (d, J = 12.3 Hz, 1H), 1.99 (t, J = 10.6 Hz, 1H), 1.88 (t, J = 10.6 Hz, 1H), 1.48 (q, J = 12.2 Hz, 1H). ^13^C NMR (101 MHz, cd3od) δ 158.95, 156.13, 150.07, 146.82, 138.79, 136.68, 133.40, 129.54, 129.46, 129.09, 128.79, 128.49, 116.58, 116.01, 113.32, 102.23, 70.95, 63.33, 60.94, 49.85, 46.14, 43.13, 41.87, 38.42. MS(ES+) m/z 470.2 [M+ H]^+^.

##### (S,S)-59-3-allyl-2-{[(3S,5S)-5-(4-(benzyloxy)phenyl)-1-methylpiperidin-3-yl]amino}I-3,5-dihydro-4H-pyrrolo[3,2-d]pyrimidin-4-one

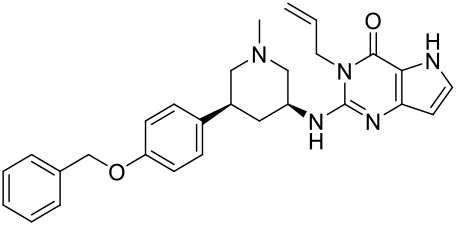

(*S,S*)-59 was prepared according to the procedure described in the literature.^14^ 5.6 mg of white solid were obtained. Rf (10% MeOH in DCM) = 0.67. ^1^H NMR (400 MHz, cd3od) δ 7.45 – 7.39 (m, 2H), 7.35 (tt, J = 6.5, 1.0 Hz, 2H), 7.35 – 7.24 (m, 1H), 7.29 – 7.13 (m, 3H), 6.98 – 6.89 (m, 2H), 6.19 – 6.10 (m, 1H), 5.99 – 5.85 (m, 1H), 5.18 (dq, J = 10.3, 1.5 Hz, 1H), 5.05 (s, 3H), 4.75 (dq, J = 3.6, 1.7 Hz, 2H), 4.40 – 4.27 (m, 1H), 3.27 (dd, J = 10.3, 3.8 Hz, 1H), 2.99 – 2.89 (m, 2H), 2.37 (s, 3H), 2.19 (d, J = 12.3 Hz, 1H), 2.07 (t, J = 11.8 Hz, 1H), 1.95 (t, J = 10.8 Hz, 1H), 1.53 (q, J = 12.2 Hz, 1H). 13C NMR (176 MHz, cd3od) δ 159.04, 156.16, 150.09, 146.82, 138.84, 136.59, 133.40, 129.55, 129.48, 129.11, 128.82, 128.51, 116.55, 116.06, 113.33, 102.21, 70.99, 63.22, 60.82, 49.80, 46.06, 43.12, 41.85, 38.36. MS(ES+) m/z 470.2 [M+ H]^+^.

### TR-FRET

The TR-FRET assay protocol was developed and adapted from a previously reported protocol.^17^ Briefly, the assay was run using white, low-volume, flat-bottom, nonbinding, 384-well microplates (Greiner, 784904) containing a total volume of 10 μL per well. The buffer was made of 1X PBS pH 7.0, 0.005% Tween 20, and 2 mM DTT. Lance Europium (Eu)-W1024 Streptavidin conjugate (2nM) and LANCE Ultra ULightTM-anti-6x-His antibody (10 nM) were used as donor and acceptor fluorophores associated with the tracer ligand and protein respectively. Final assay concentrations of 40 nM 6X histidine tagged SETDB1 (residues 195-403, N-terminal tag) and 40 nM of H3K9Me2K14Ac-biotin (aa 1-19) were used for compound testing. A 10 point, three-fold serial dilution of each compound was tested as the primary hit validation. Assay components were added to an assay plate using a Multidrop Combi (ThermoFisher). After addition, the plates were sealed with clear covers, mixed gently on a shaker for 1 min, centrifuged at 1000x g for 2 minutes, and allowed to equilibrate for 1 hour in the dark. Plates were read with an EnVision 2103 Multilabel Plate Reader (PerkinElmer) using and excitation filter at 320 nm and emission filters at 615 and 655 nm. Emission signals (615 and 665 nm) were measured simultaneously using a dual mirror D400/D630 (using a 100-microsecond delay). TR-FRET output sign was expressed as emission ratios of acceptor/donor (665/615 nm) counts. Percent inhibition was calculated on a scale of 0% (i.e. activity with DMSO vehicle only) to 100% (100 μM H3K9Me2K14Ac) using two full columns of control wells on each plate. The data was fitted with a four-parameter nonlinear regression analysis using ScreenAble to determine IC50 values, plotted on GraphPad for visualization, and reported as an average of at least three technical replicates ± SEM.

### DSF

DSF assays were performed using AB ViiA 7 Real-Time PCR System. The buffer was made of 20 mM HEPES, 200 mM NaCl, Ph 7.5. Experiments were carried out using 8 μL of SETDB1-3TD (residues 196-403, N-terminal His tag) and 20X concentration Sypro Orange dye (5000X DMSO stock, ThermoFisher) and 2 uL of ligand for a final concentration of 20 μM of protein and 200 μM of ligand with 1.1% of DMSO. Plates were uncubated for 15 min before running the temperature gradient (1 °C/min, from 25 to 90 °C). Values are presented as an average of at least three replicates ± SEM calculated using the Boltzmann method from the Protein Thermal Shift software (ThermoFisher).

#### Surface Plasmon Resonance

SPR experiments were carried out at 25 °C on a Biacore 8K SPR system (GE Healthcare). Series S SA sensor chips (GE Healthcare) were used in all experiments. All experiments were run in an identical buffer made of 20 mM Tris/HCl [pH 7.0], 150 mM NaCl, and 0.005% Tween20 (vol/vol), at 1% DMSO. Biotinylated SETDB1-TTD was first immobilized on the SA chip (injection concentration: 10 μg/mL, contact time: 60 s, flow rate: 10 μL/min) to achieve an optical density of approximately 5000-6000 RU. Next, varying concentrations of test compounds were flowed over the immobilized protein in a dose-response fashion (contact time: 120 s, dissociation time: 300 s, flow rate: 30 μL/min) using a multi-cycle kinetics protocol. The resulting data was analyzed using the Biacore Evaluation Software (GE Healthcare) and plotted on GraphPad for visualization. SPR Kd values are reported as an average of at least three technical replicates ± SEM.

#### Cell culture and transfection

Human breast cancer cell line MDA-MB-231, human colorectal adenocarcinoma cell DLD-1 and human immortalized kidney cell lines HEK293T were purchased from ATCC and cultured in DMEM medium supplemented with 10% FBS, 100U penicillin and 100mg/mL streptomycin. Cell transfection was performed using polyethylenimine (PEI) (23866-1, Polysciences, Inc) according to manufacturer instructions.

#### Plasmid construction

pcDNA3-HA-Akt1 was constructed by cloning corresponding PCR fragments into pcDNA3-HA vector by BamHI and SalI sites.

Akt1-BamHI-Forward:

5′-CATGGATCCAGCGACGTGGCTATTGTG-3′

Akt1-SalI-Reverse:

5′-GCATGTCGACTCAGGCCGTGCCGCTGGC-3′

#### Immunoblot and Immunoprecipitations Analyses

Cells were lysed in EBC buffer (50 mM Tris pH 7.5, 120 mM NaCl, 0.5% NP-40) or Triton X-100 buffer (50 mM Tris, pH 7.5, 150 mM NaCl, 1% Triton X-100) supplemented with protease inhibitor cocktail and phosphatase inhibitor cocktail. The protein concentrations of whole cell lysates were measured by NanoDrop OneC using the Bio-Rad protein assay reagent. Equal amounts of whole cell lysates were loaded by SDS-PAGE and immunoblotted with indicated antibodies. For immunoprecipitations analysis, 1 mg total lysates were incubated with the anti-HA agarose beads (A-2095, Sigma) for 3-4 hr at 4°C. The recovered immuno-complexes were washed three times with NETN buffer (20 mM Tris, pH 8.0, 100 mM NaCl, 1 mM EDTA and 0.5% NP-40) before being resolved by SDS-PAGE and immunoblotted with indicated antibodies.

#### Antibodies

All antibodies were used at a 1:1,000 dilution in TBST buffer with 5% non-fat milk for western blotting unless specified. Anti-HA-Tag antibody (3724), anti-Tri-Methyl Lysine motif antibody (14680) and Anti-phospho-Akt-Thr308 antibody (9275) were obtained from Cell Signaling Technology.

#### Cell Proliferation Assays

5,000 indicated cells were seeded in each well of 96-well plates followed by treatment with indicated doses of compounds for 72 hrs before cell proliferation analysis by Cell Titer-Glo Kit (G9241, Promega) according to manufacture instructions. Corresponding luminescent signals were measured using the BioTek Cytation 5 Cell Imaging reader.

## CONCLUSION

SETDB1 is a large multi-domain protein involved in cancer development through several mechanisms including Akt1 activation. In this study, we confirmed that SETDB1 is able to methylate K64 of Akt1. We showed that a 3TD-lacking, catalytically active, SETDB1 construct (SETDB1-S) only displayed a weak ability to install the methyl mark on Akt1-K64 *in vitro*, compared to SETDB1-FL, suggesting that the 3TD plays a key role in the methyltransferase function of SETDB1. We then demonstrated that the TD2 ligand, (*R,R*)-59, is able to increase the methyltransferase activity of SETDB1 both *in vitro* and in cells. Binding of (*R,R*)-59 to the TD2 stimulates the methyl transfer from SAM to K64 of Akt1 potentially through an allosteric interaction between the 3TD and the SET domain of SETDB1. Moreover, after treatment with (*R,R*)-59, SETDB1-mediated methylation of Akt1 in cells and subsequent T308 phosphorylation results in Akt1 activation. Akt1 activation is a known carcinogenic pathway and, as expected, (*R,R*)-59 treatment resulted in increased cell proliferation in breast cancer cells where SETDB1 is commonly overexpressed. Overall, these results demonstrate the ability of (*R,R*)-59 to activate SETDB1 methyltransferase activity both *in vitro* and in cells, suggesting that (*R,R*)-59 is the first SETDB1 small molecule positive allosteric modulator. Further studies to evaluate SETDB1-mediated histone H3 methylation and the exact mechanism of action of (*R,R*)-59 are being further pursued to gain a better understanding of the relationship between the 3TD and SET domain of SETDB1. Given the interest in converting protein ligands into targeted degradation agents, our finding that (*R,R*)-59 acts as a positive activator toward SETDB1 provides an important caveat when starting from ligands of the 3TD of SETDB1.

## ASSOCIATED CONTENT

### Supporting Information

Figure S1: TR-FRET evaluation of (R,R)-59 and (S,S)-59 for the displacement of H3K9Me2K14Ac from the Triple Tudor Domain of SETDB1; Figure S2: SPR evaluation of (*R,R*)-59 for the 3TD of SETDB1; Figure S3: Kinetic evaluation of the influence of Akt1-K64 on the enzymatic activity of SETDB1 with and without (*R,R*)-59; ^1^H, ^13^C NMR and LC-MS characterization of (*R,R*)-59 and (*S,S*)-59 (PDF)

## ACKNOWLEDGMENT

This work was supported by the PhRMA Foundation (Drug Discovery postdoctoral fellowship) to M.U. and the National Institute of General Medical Sciences, NIH (grant R35GM139514) to S.V.F. The authors thank P. Buttery and R. Johnson for reviewing the experimental chemistry and biology data. The authors thank B. Hardy for assembly of the screening plate for TR-FRET. The Structural Genomics Consortium is a registered charity (no: 1097737) that receives funds from Bayer AG, Boehringer Ingelheim, Bristol Myers Squibb, Genentech, Genome Canada through Ontario Genomics Institute [OGI-196], EU/EFPIA/OICR/McGill/KTH/Diamond Innovative Medicines Initiative 2 Joint Undertaking [EUbOPEN grant 875510], Janssen, Merck KGaA (aka EMD in Canada and US), Pfizer and Takeda. Our abstract figure was generated by biorender.com.

