## Supporting information for "SETDB1 Triple Tudor Domain Ligand, (*R,R*)-59, Promotes Methylation of Akt1 in Cells"

### 1. Supplementary Figures

#### 1.1. Figure S1

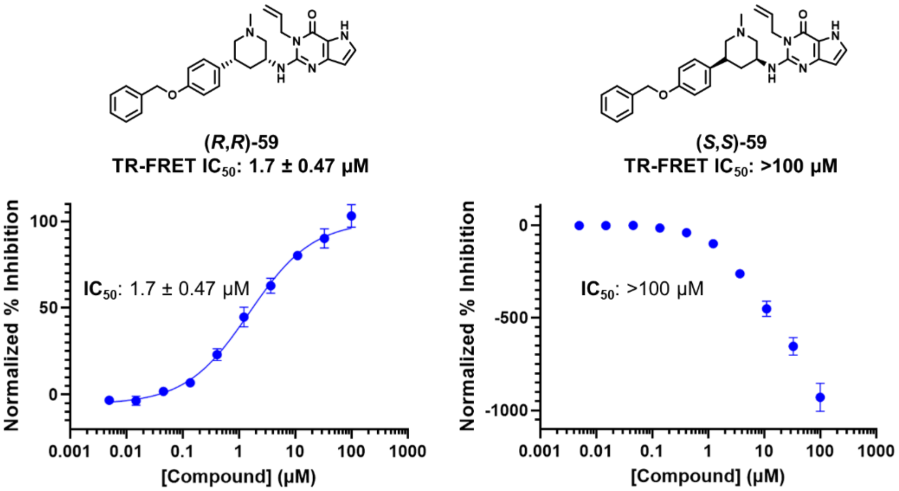

**Figure S1.** TR-FRET evaluation of (*R*,*R*)-59 and (*S*,*S*)-59 for the displacement of H3K9Me2K14Ac from the Triple Tudor Domain of SETDB1.

#### 1.2. Figure S2

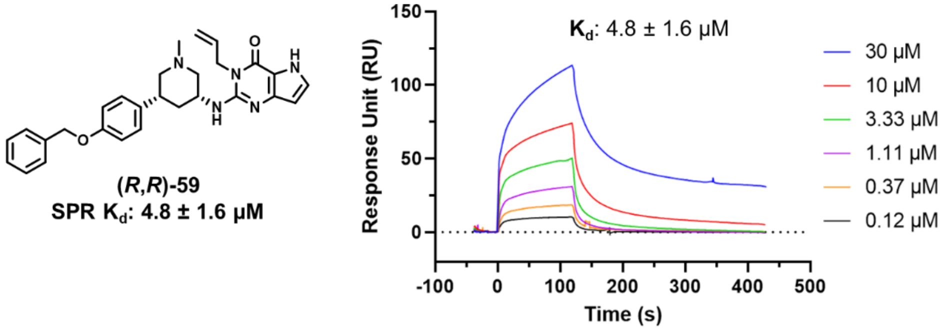

**Figure S2.** SPR evaluation of (*R*,*R*)-59 for the 3TD of SETDB1.

#### 1.3. Figure S3

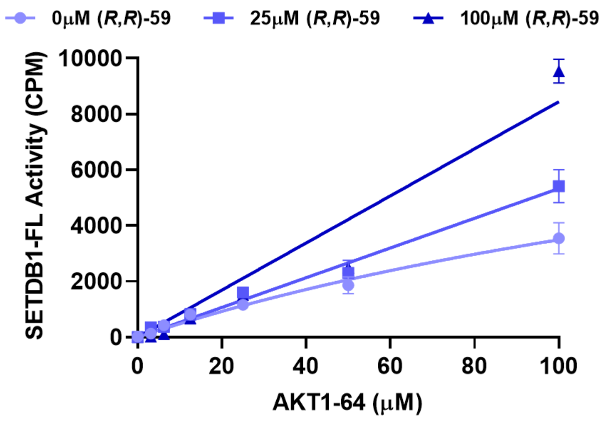

**Figure S3.** Kinetic evaluation of the influence of Akt1-K64 on the enzymatic activity of SETDB1 with and without (*R*,*R*)-59.

#### 2. NMR and LC-MS Data

##### 2.1. $^{1}$H NMR and $^{13}$C NMR of (*R*,*R*)-59

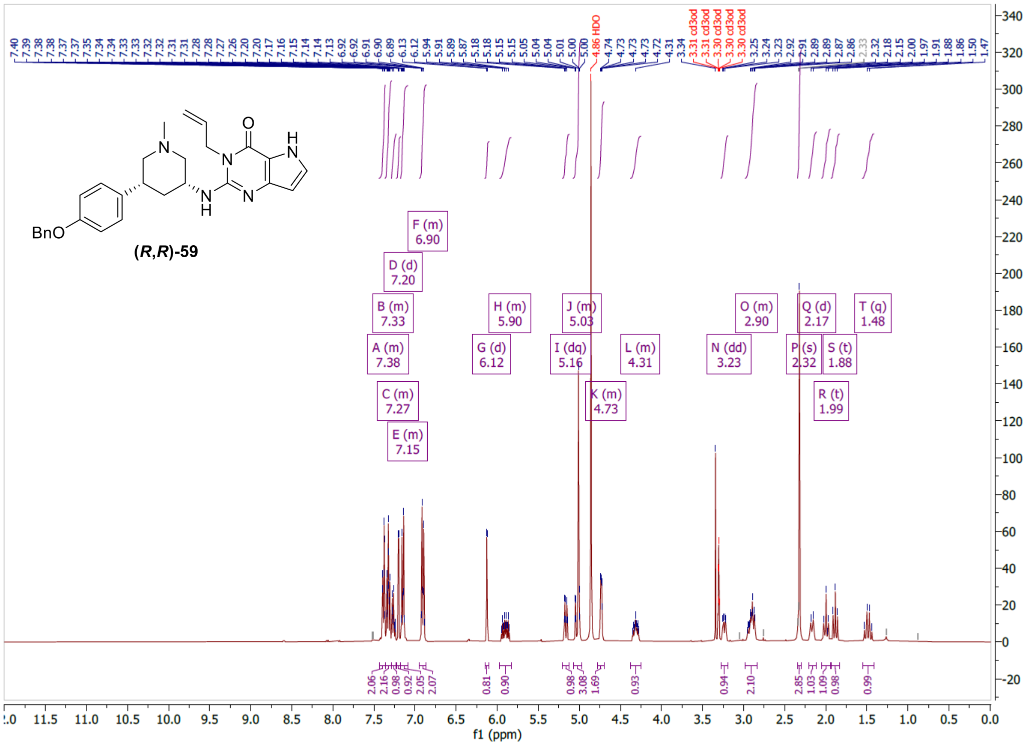

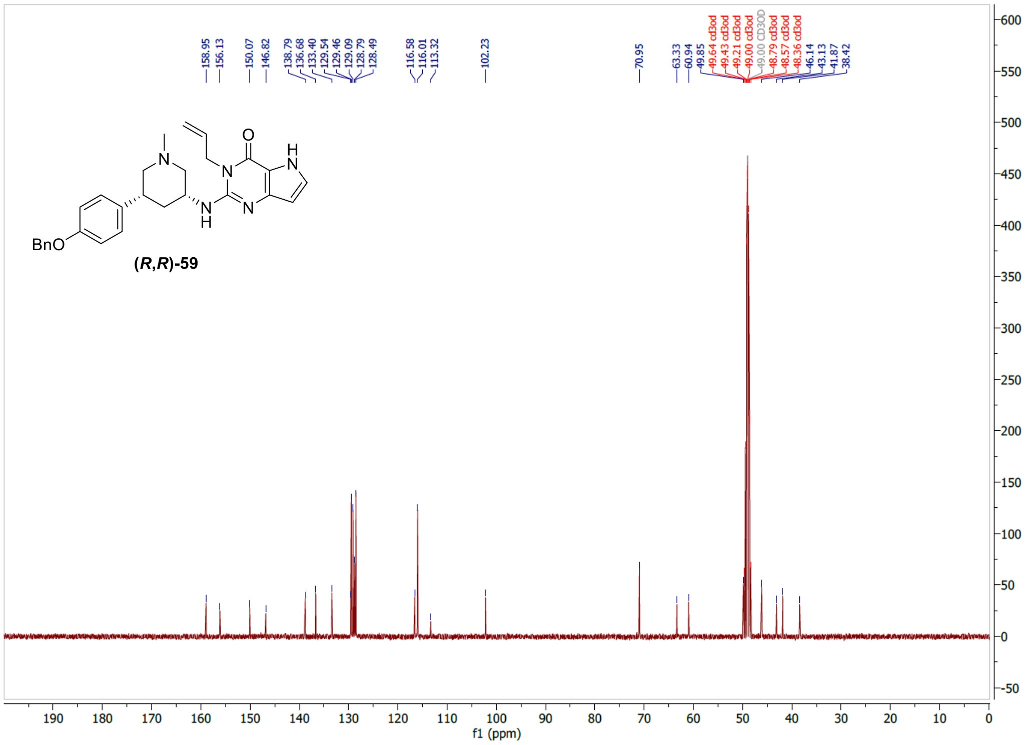

#### 2.2. LC-MS data for (*R*,*R*)-59

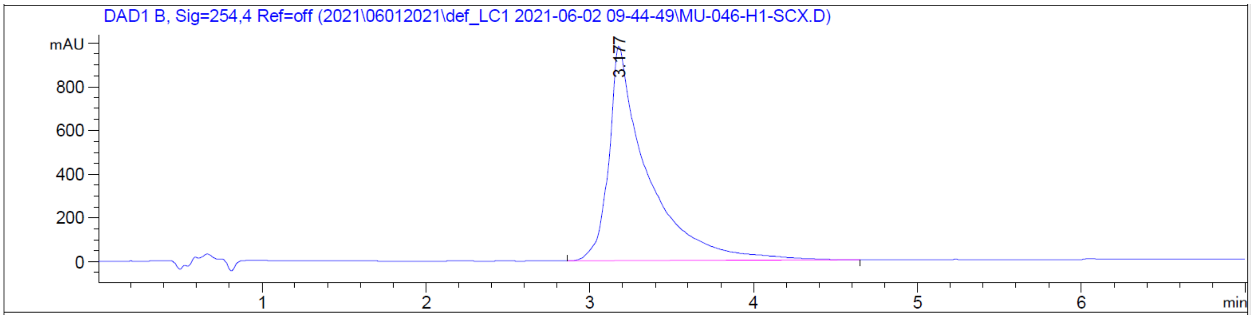

#### 2.3. [1]H NMR and [13]C NMR of (*S*,*S*)-59

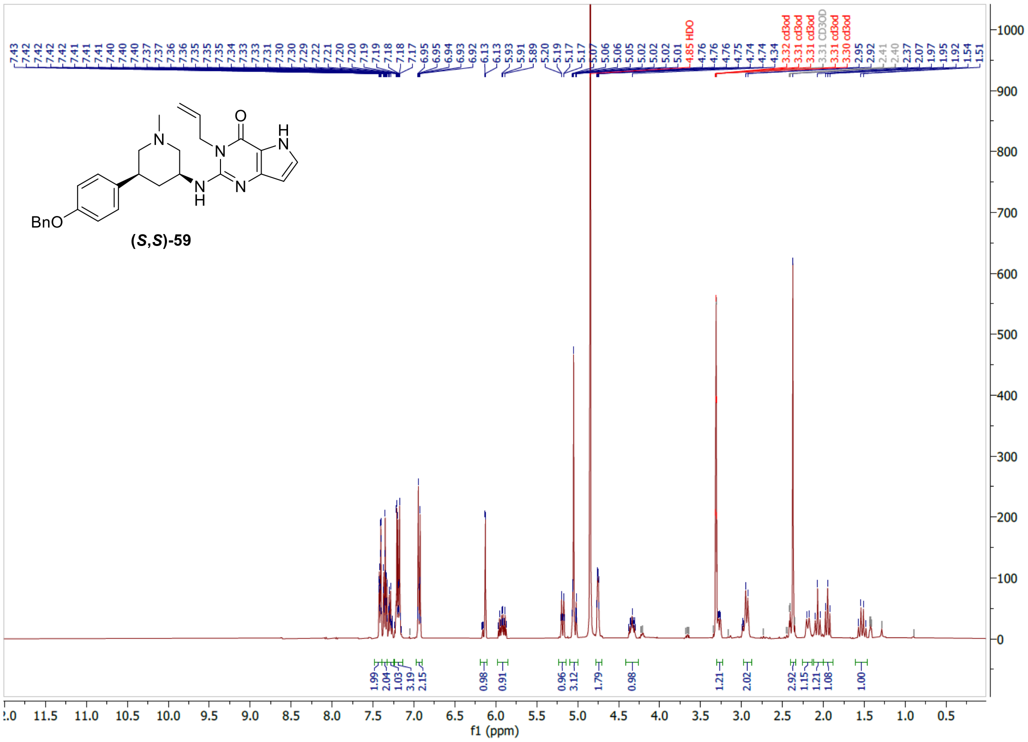

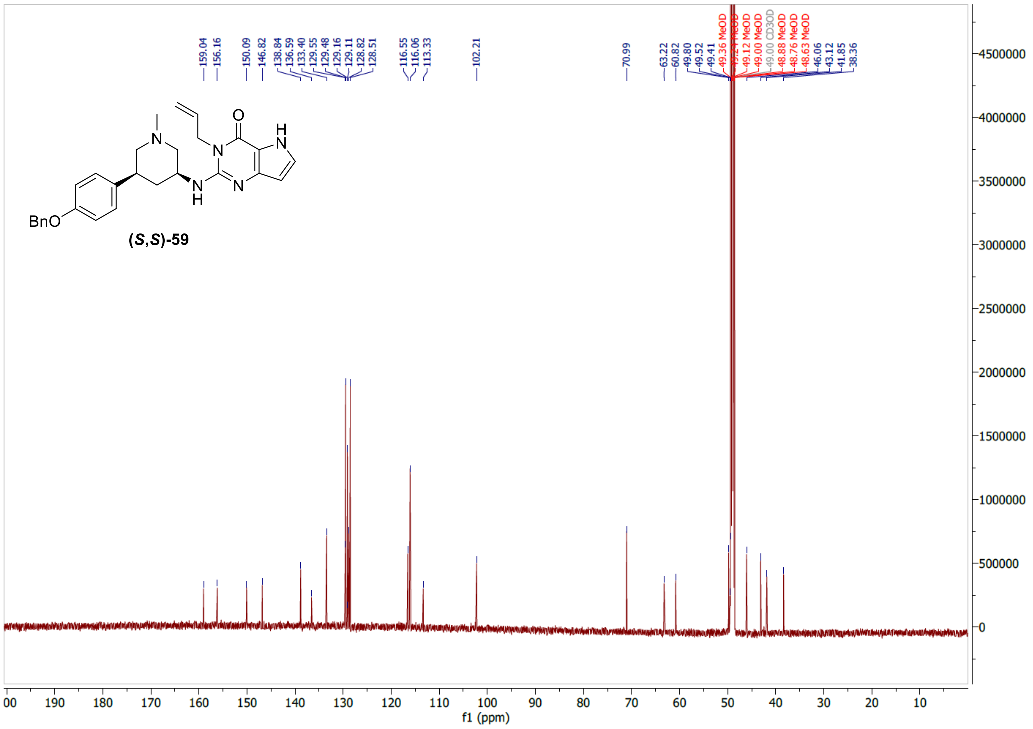

#### 2.4. LC-MS data for (*S*,*S*)-59

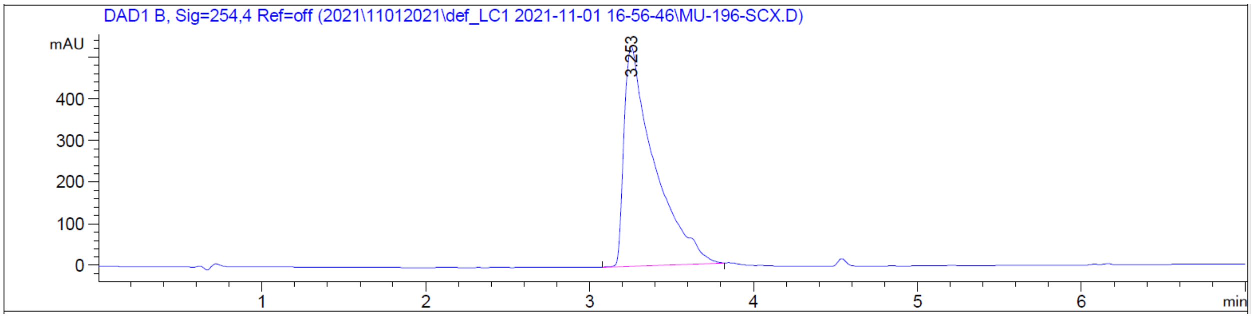
